# Aperiodic pupil fluctuations at rest predict orienting of visual attention

**DOI:** 10.1101/2024.04.26.591420

**Authors:** Rongwei Wang, Jianrong Jia

## Abstract

The aperiodic exponent features signals in various neuroimaging modalities and has been linked to the excitation/inhibition balance of the neural system. Leveraging the rich temporal dynamics of resting-state pupil fluctuations, the present study investigated the association between the aperiodic exponent of pupil fluctuations and the neural excitation/inhibition balance in attentional processing. In separate phases, we recorded participants’ pupil size during resting state and assessed their attentional orienting using the Posner cueing tasks with different cue validities (i.e., 100% and 50%). We found significant correlations between the aperiodic exponent of resting pupil fluctuations and both the microsaccadic and behavioral cueing effects. Critically, this relationship was particularly evident in the 50% cue-validity condition rather than in the 100% cue-validity condition. The microsaccadic responses mediated the association between the aperiodic exponent and the behavioral response. Further analysis showed that the aperiodic exponent of pupil fluctuations predicted the self-rated hyperactivity/impulsivity trait across individuals, suggesting its potential as a marker of attentional deficits. These findings highlight the rich information contained in pupil fluctuations and provide a novel approach to assessing the neural excitation/inhibition balance in attentional processing.

## Introduction

Brain activity at different spatiotemporal scales often shows a mixture of periodic and aperiodic fluctuations. While the periodic components produce peaks in power spectra, the aperiodic component manifests as a background spectral trend that decreases power with increasing frequency. This aperiodicity has been shown in the fluctuations of electrocorticography (ECoG, Manning et al., 2009), electroencephalogram (EEG, Waschke et al., 2021), magnetoencephalography (MEG, Dehghani et al., 2010), and functional magnetic resonance imaging (fMRI, He, 2011) signals, as well as in animal (Jones et al., 2023) and human (Palva et al., 2013) behavioral recordings. Intriguingly, while the slopes of the power spectrum are different for different neuroimaging modalities, their aperiodic fluctuations are strongly related across the modalities (Jones et al., 2023), indicating a general neural mechanism underlying these fluctuations. Evidence has suggested that the slope of the power spectrum (i.e., the aperiodic exponent) tracks the ratio of excitation and inhibition in the underlying neuronal circuits (Gao et al., 2017). For instance, causal evidence has been found where the anesthetic drug propofol, which enhances neural inhibition, increases the slope of the power spectrum, while the anesthetic drug ketamine, which enhances neural excitation, decreases the slope of the power spectrum (Waschke et al., 2021). In summary, the aperiodic fluctuations of the various neural signals are suggested to be reflections of the excitation/inhibition (E/I) balance in the associated neural system.

Aperiodic fluctuations of EEG signal were found to be strongly associated with attentional processing. Pre-stimulus aperiodic fluctuations predicted subjective awareness of auditory and visual stimuli (Cunningham et al., 2023), while post-stimulus aperiodic exponent was associated with the spatial distribution of attention (Pietrelli et al., 2022). EEG aperiodic exponent during resting state predicted individuals’ temporal resolution of perception (Deodato & Melcher, 2023). These studies suggest that aperiodic fluctuations in the EEG signal reflect the neural E/I balance in attentional processing (Cunningham et al., 2023; Deodato & Melcher, 2023; Pietrelli et al., 2022). Changes in the aperiodic exponent of the EEG signal have also been found to be associated with attention-deficit/hyperactivity disorder (ADHD). For example, Robertson et al. (2019) reported that the spectral slopes were steeper in preschool-age children with ADHD. Consistently, children with ADHD have been found to show a greater theta-beta ratio (Arns et al., 2013), which also implies a steeper spectral slope (Finley et al., 2022). However, other researchers have found flatter spectral slopes in school-age and adolescent ADHD samples (Ostlund et al., 2021). Karalunas et al. (2022) reconciled this conflict by showing that infants with a family history of ADHD had steeper spectral slopes, whereas adolescents diagnosed with ADHD had flatter slopes, suggesting that the trend reverses across age.

Like neural activity and behavioral responses, pupil size fluctuates dynamically over time (Reimer et al., 2016), even at rest. Extensive evidence has suggested that raw pupil size tracks the degree of attention and the state of arousal (Contadini-Wright et al., 2023; Joshi & Gold, 2020), but direct examinations of the pupil fluctuations, the aperiodic exponent of pupil fluctuations in particular, are lacking. There is indirect evidence to imply the association between pupil fluctuations and the neural E/I balance. Pfeffer et al. (2022) recently showed a negative correlation between pupil size and the aperiodic exponent of MEG signal. Critically, pupil fluctuations, such as the hippus (oscillations below 1 Hz), reflect neural excitability better than raw pupil size (Binda & Lunghi, 2017; Pomè et al., 2020). These findings inspired the present study to investigate whether the aperiodic exponent of pupil fluctuations could also reflect neural E/I balance, just like neural signals in other neuroimaging modalities. Furthermore, being controlled by the locus coeruleus-norepinephrine (LC-NE) and the basal forebrain-acetylcholine (BF-ACh) systems (Thiele & Bellgrove, 2018), the pupil fluctuations may specifically be associated with the neural E/I balance in attentional processing, which is also controlled by these two neurotransmitter systems.

To investigate the association between the aperiodic pupil fluctuations and the neural E/I balance in attentional processing, we examined the correlation between the aperiodic exponent of the resting pupil and the attentional orienting across individuals. Pupil size was recorded during the resting state, while participants’ behavioral and microsaccadic responses to exogenous attention were measured using two Posner cueing tasks of different cue validities (100% and 50%) (Fig. 1A). Results showed significant correlations between the aperiodic exponent of resting pupil and both the behavioral and microsaccadic cueing effects when cue validity was 50% rather than 100%. The microsaccadic responses mediated the association between the aperiodic exponent and the behavioral response. To validate the association between the aperiodic exponent of the resting pupil and the attentional orienting, we further measured participants’ self-rated ADHD symptoms using the Adult Self-Rating Scale (ASRS, Kessler et al., 2005). Replicating the findings in the attentional tasks, the aperiodic exponent was predictive of self-rated hyperactivity/impulsivity. Together, the present study demonstrates for the first time that aperiodic pupil fluctuations predict the orienting of visual attention and suggests an association between aperiodic pupil fluctuations and neural E/I balance in attentional processing.

**Fig. 1.**
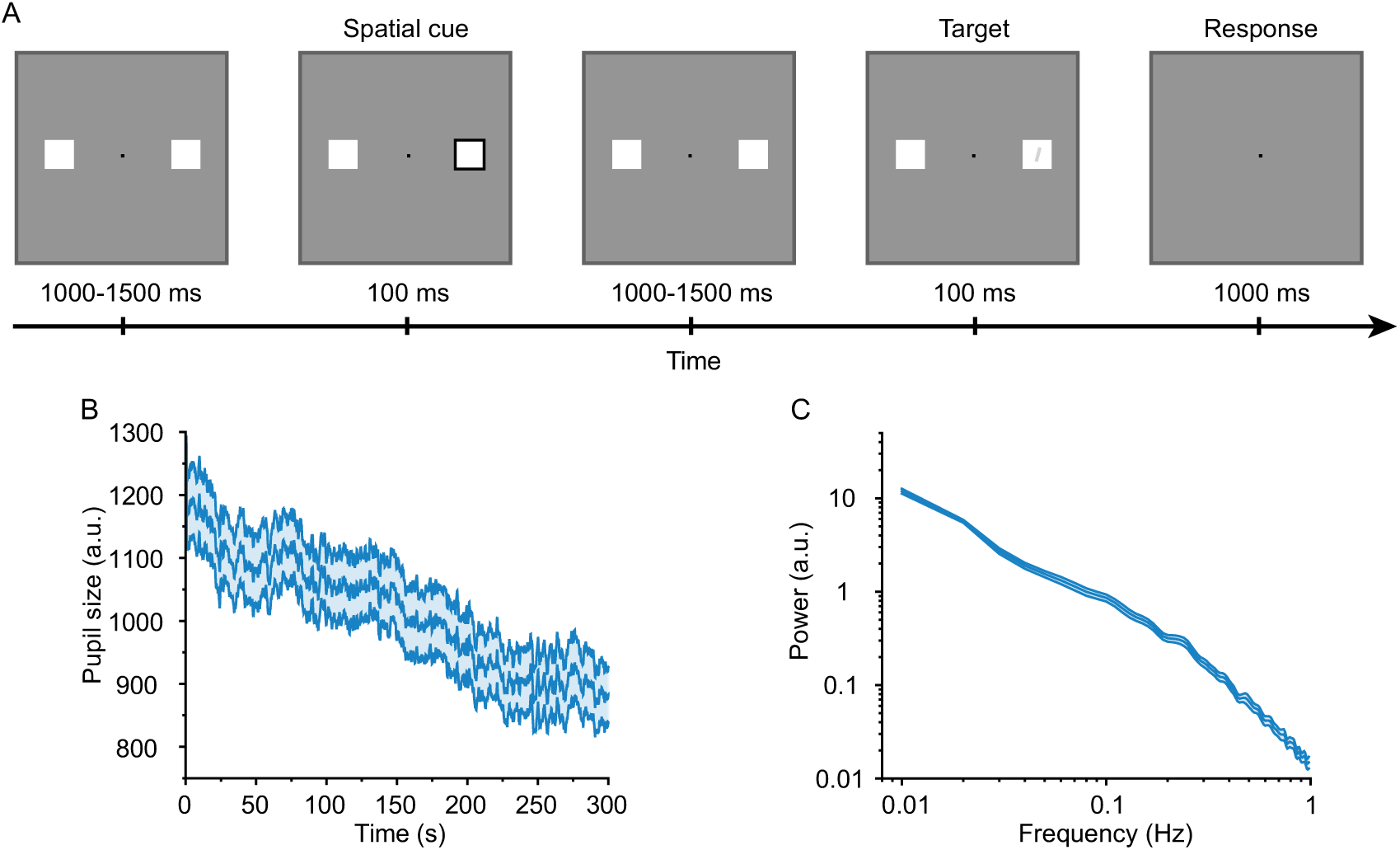
The trial structure of the experimental task and the pupil fluctuations in the 300-s resting state. (A) The trial structure for a valid-cue condition of the task. (B) The fluctuations of the raw pupil size during the 300-s resting state. (C) The PSD of the resting-state pupil fluctuations was shown in the double logarithmic coordinates. The shaded areas in B and C indicate the standard error of the mean (SEM).

## Methods

### Participants

The sample size was calculated using G*Power software (version 3.1.9.6), which suggested that a minimum of 69 participants was required for a reliable correlational study. This calculation was based on a power of 0.8, an alpha error rate of 0.05, and a correlation coefficient of 0.33 derived from a similar study (Ghirardi et al., 2019). In the present study, 70 adult participants (33 males, 19-25 years old) were recruited. All participants had normal or corrected-to-normal vision and had no known neurological or visual disorders. The research protocols were approved by the Ethics Committee of the Affiliated Hospital of Hangzhou Normal University, approval number 2022(E2)-KS-118. Written informed consents were obtained from all participants prior to the experiment.

### Apparatus and tools

The visual stimuli were presented on a CRT monitor (resolution: 1024×768; refresh rate: 85 Hz). The MATLAB (The MathWorks) and the Psychophysics Toolbox (Brainard, 1997) were used to present the stimuli and record the behavioral responses. The experiment took place in a dimly lit and sound-proof room. The participants comfortably seated at a viewing distance of about 70 cm, with their heads stabilized on a chin rest.

### Stimuli and procedure

The experiment consisted of a spatial cueing task phase and a resting-state eye-movement recording phase. The spatial cueing task was similar to that in our previous study on attentional orienting (Jia et al., 2017; Lv et al., 2022). Fig. 1A shows the events that occurred in a typical trial. At the beginning of each trial, a central fixation dot (0.16°×0.16°; 0 cd/m^2^), flanked by two square place holders (4°×4°; 104 cd/m^2^; distance to the fixation dot: 6°), was presented on a gray background (23 cd/m^2^) for 1000-1500 ms. Then, the border of one of the placeholders suddenly dimmed to 0 cd/m^2^ in luminance for 100 ms, serving as exogenous cueing on the corresponding spatial location. Following an inter-stimulus interval of 1000-1500 ms, a bar (size: 1.30°×0.13°; luminance: 67 cd/m^2^) tilted left or right by 2° appeared in one of the placeholders for 100 ms as a probe. Participants were instructed to report the tilt orientation of the probe as accurately as possible by pressing one of two buttons within 1100 ms. The participants were required to maintain fixation throughout the whole trial. The participants completed two blocks of the spatial cueing task, with the validity of the spatial cue being 100% and 50%, respectively. The assignment of the validity to each block was counterbalanced across participants, and participants were informed about the cue validity before starting each block. Each block contained 200 trials, taking about 20 minutes to complete. Participants completed 15 practical trials at the beginning of each block.

After the spatial cueing task, participants underwent a resting-state phase, in which their eye movement was recorded. During this phase, participants were required to look at the fixation point (0.16°×0.16°; 0 cd/m^2^) in the center of the gray screen (23 cd/m^2^) for 300 s and were told to minimize blinking.

### Eye movement recording

Eye movements were recorded binocularly at 1000 Hz with an EyeLink 1000 eye tracker (SR Research, Ottawa) during the whole experiment. To maintain good tracking accuracy, each participant used a chin rest. At the beginning of each experimental block, the eye tracker was calibrated with a standard 9-point calibration procedure.

### Microsaccade detection in the spatial cueing task

Microsaccades were detected with an improved version (Engbert & Mergenthaler, 2006) of the algorithm originally proposed by Engbert and Kliegl (2003). Horizontal and vertical eye positions were mapped onto a velocity space, to which a 5 standard deviation threshold was applied to detect saccades. For a detected saccade to be counted as a microsaccade, three criteria must meet: 1) a temporal overlap between the two eyes, 2) a minimum duration of 6 ms, and 3) an amplitude below 1°. Trials that contained saccades larger than 1° were excluded from the analysis. This data cleansing procedure excluded 8% and 10% of the trials from the 100% and 50% cue-validity conditions, respectively.

### Microsaccade rate analysis

A rectangular moving window of 100 ms (stepped in 1 ms) was used to examine the microsaccade rate in a time window of 100 ms before and 1000 ms after the attention cue onset (Engbert & Kliegl, 2003). Given that a microsaccade could land in the same or opposite hemifield as the cue, we referred to these as “congruent” and “incongruent” microsaccades, respectively. The cueing effect refers to the rate difference between congruent and incongruent microsaccades.

### Resting-state pupillometry data processing

The PuPl toolbox (Kinley & Levy, 2021) was used for pre-processing of the raw pupil size data, which took the following three steps:

1. Raw pupil size data cleaning. The samples in which pupil size was changing at an implausibly high rate (i.e., where the “dilation speed” is too high) and samples that were small “islands” surrounded by missing data were recognized as outliers and deleted from the recordings (Kret & Sjak-Shie, 2019). The threshold of the dilation speed was set to be the median speed plus 3 times the median absolute deviation. The “islands” were defined as samples with a duration of less than 30 ms and were surrounded by missing data exceeding 25 ms.
2. Blink identification. Blinks were identified with the pupillometry noise method (Hershman et al., 2018). The identified blink samples were deleted from the recordings along with their adjacent data within 50 ms prior to and 150 ms after the identified blinks.
3. Data interpolation, smoothing, and downsampling. The gaps in pupil size data resulting from data cleaning and blink removal were filled in using linear interpolation. High-frequency noises in the pupil size data were removed by data smoothing using a moving mean filter with a 150-ms length Hanning window (Zhao et al., 2019). Finally, the data were downsampled from 1000 Hz to 100 Hz and averaged across two eyes.

### Spectral slope and power analysis

The mean value was first subtracted from the pre-processed 300 s pupil data. Then, the power spectral density (PSD) of the pupil data was calculated using Welch’s method with a window length of 5000 data points and 50% overlapping windows. The “Fitting Oscillations and One-Over-f” (FOOOF) toolbox was used to calculate the aperiodic exponent (Donoghue et al., 2020) of the PSDs, which uses a spectral parameterization algorithm to decompose the power spectrum into periodic and aperiodic components via an iterative model fitting process. Considering that the fluctuations of pupil size are mainly concentrated between 0-1 Hz (Pomè et al., 2020; i.e., pupillary hippus; Turnbull et al., 2017) and that the spectral plateau above 1 Hz could disrupt the estimation of aperiodic exponent (Gerster et al., 2022), we extracted aperiodic exponents from the 0-1 Hz frequency range of each power spectrum (aperiodic_mode = ‘fixed’, max_n_peaks = ‘inf’, default settings otherwise). The ‘fixed’ setting was used given that we did not anticipate a “knee” in the power spectrum. This assumption was supported by visual inspection of each PSD after spectral parameterization via FOOOF. The value of the aperiodic exponent was high for the steeper spectrum and low for the flatter spectrum.

### ADHD-like symptoms measurement

ADHD-like symptoms were measured using the World Health Organization Adult ADHD Self-Report Scale (ASRS; Kessler et al., 2005), an eighteen-item measure of ADHD-like symptoms that can be administered verbally or through survey. The self-rated “inattention” and “hyperactivity/impulsivity” symptoms were each measured with nine items of the ASRS. Participants select how frequently each statement has been true in the past 6 months on a five-point scale from 0 (never) to 4 (very often). Higher scores indicate more severe self-rated ADHD-like traits.

### Statistical analysis

Pearson’s correlation was used to examine the correlation between different variables. As significant correlations were found between the aperiodic exponent and the microsaccadic cueing effect and between the aperiodic exponent and the behavioral cueing effect, a formal participant-wise mediation model was implemented. The indirect path from the aperiodic exponent to the microsaccadic cueing effect (*a*) and the microsaccadic cueing effect to the behavioral cueing effect (*b*) was computed, as was the direct path from the aperiodic exponent to the behavioral cueing effect (*c’*). The indirect path and mediation were computed as the product of *a* × *b*, with a resulting 95% confidence interval for each indirect path from 1,000 bootstrapped samples. The total, direct, and indirect effects were tested for significance at P < 0.05. The mediation analysis was performed using JASP (version 0.18.0) software.

## Results

### The aperiodic exponent of resting pupil fluctuations was associated with the behavioral cueing effect

We first validated the pupil size recordings by examining the characteristics of pupil fluctuations. Fig. 1B shows that the pupil size in the resting state gradually decreased over time during the 300-s recording phase. The power spectral density (PSD) of the pupil size fluctuations (Fig. 1C) followed a 1/f power law distribution, with the power being higher at lower frequencies and decreasing rapidly with increasing frequency. The decrease in power was roughly linear in the double logarithmic coordinates (Fig. 1C). We used the FOOOF algorithm (Donoghue et al., 2020) to separate the aperiodic and oscillatory components of the pupil spectrum. Next, our analysis focused on the relationship between the aperiodic exponent of pupil fluctuations and the behavioral performance in the attention task.

We calculated the correlations between the aperiodic exponent and both the response accuracies and cueing effect. The correlations between the aperiodic exponent and the response accuracies were not significant in either the 100% (*r* = 0.20, *P* = 0.100; Fig 2A) or the 50% (*r* = 0.18, *P* = 0.138; Fig 2B) cue-validity conditions. These results suggest that the aperiodic exponent of pupil fluctuations is not predictive of the overall accuracy in attentional tasks, either in the 100% or the 50% cue-validity conditions.

**Fig. 2.**
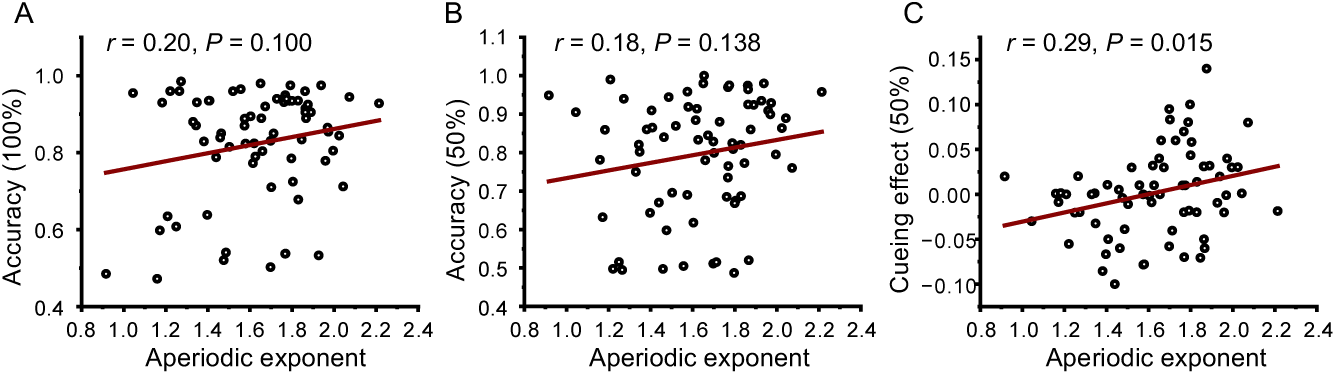
The correlations between the aperiodic exponent of pupil fluctuations and behavioral performance. (A) The correlation between the aperiodic exponent and the response accuracy in the 100% cue-validity condition. (B) The correlation between the aperiodic exponent and the response accuracy in the 50% cue-validity condition. (C) The correlation between the aperiodic exponent and the cueing effect in the 50% cue-validity condition.

Next, we calculated the correlation between the aperiodic exponent and the spatial cueing effect (i.e., the difference between the response accuracy in the cued direction and that in the uncued direction) in the 50% cue-validity condition (i.e., the cue did to predict where the probe would appear). The result showed a significant correlation (*r* = 0.29, *P* = 0.015; Fig 2C) that the larger aperiodic exponent was accompanied by the positive cueing effect. A control analysis showed that the aperiodic exponent was not significantly correlated with accuracy for either cued (*r* = 0.18, *P* = 0.139) or uncued (*r* = 0.20, *P* = 0.094) targets alone. The clear association between the aperiodic exponent and the attentional cueing effect rather than the overall accuracy suggests that the aperiodic exponent of pupil fluctuations predicts the balance of attention in the cued and uncued directions.

### The aperiodic exponent of resting pupil fluctuations was correlated with the microsaccadic IOR in the 50% cue-validity condition

Attentional effects elicited by exogenous cues are reflected by dynamic changes in the microsaccade rate (Laubrock et al., 2005; Lv et al., 2022). The microsaccade following the presentation of an exogenous cue was found to be more directed in the opposite direction of the cue, reflecting a microsaccadic inhibition of return (IOR; Galfano et al., 2004; Laubrock et al., 2005; Lv et al., 2022; Rolfs et al., 2004). We further examined the relationship between the aperiodic exponent of pupil fluctuations and the microsaccadic responses. For illustration purposes, we first divided the participants into two groups according to the median value of the aperiodic exponent. We calculated the cueing effect for the microsaccade rate by subtracting the rate of microsaccade in the uncued direction from that in the cued direction. For the 100% cue-validity condition, the cueing effect of the microsaccade rate was negative after cue presentation (Fig. 3A), replicating the classic microsaccadic IOR effect (Laubrock et al., 2005; Lv et al., 2022). Critically, the microsaccadic cueing effect was not different between the two groups of participants with high and low aperiodic exponents. To validate the results of median splitting, we extracted the microsaccadic cueing effect in 200-500 ms for all participants and calculated its correlation with the aperiodic exponent of the pupil fluctuations (Fig. 3B). We found that the aperiodic exponent was not significantly correlated with the microsaccadic cueing effect in the 100% cue-validity condition (*r* = 0.01, *P* = 0.946).

**Fig. 3.**
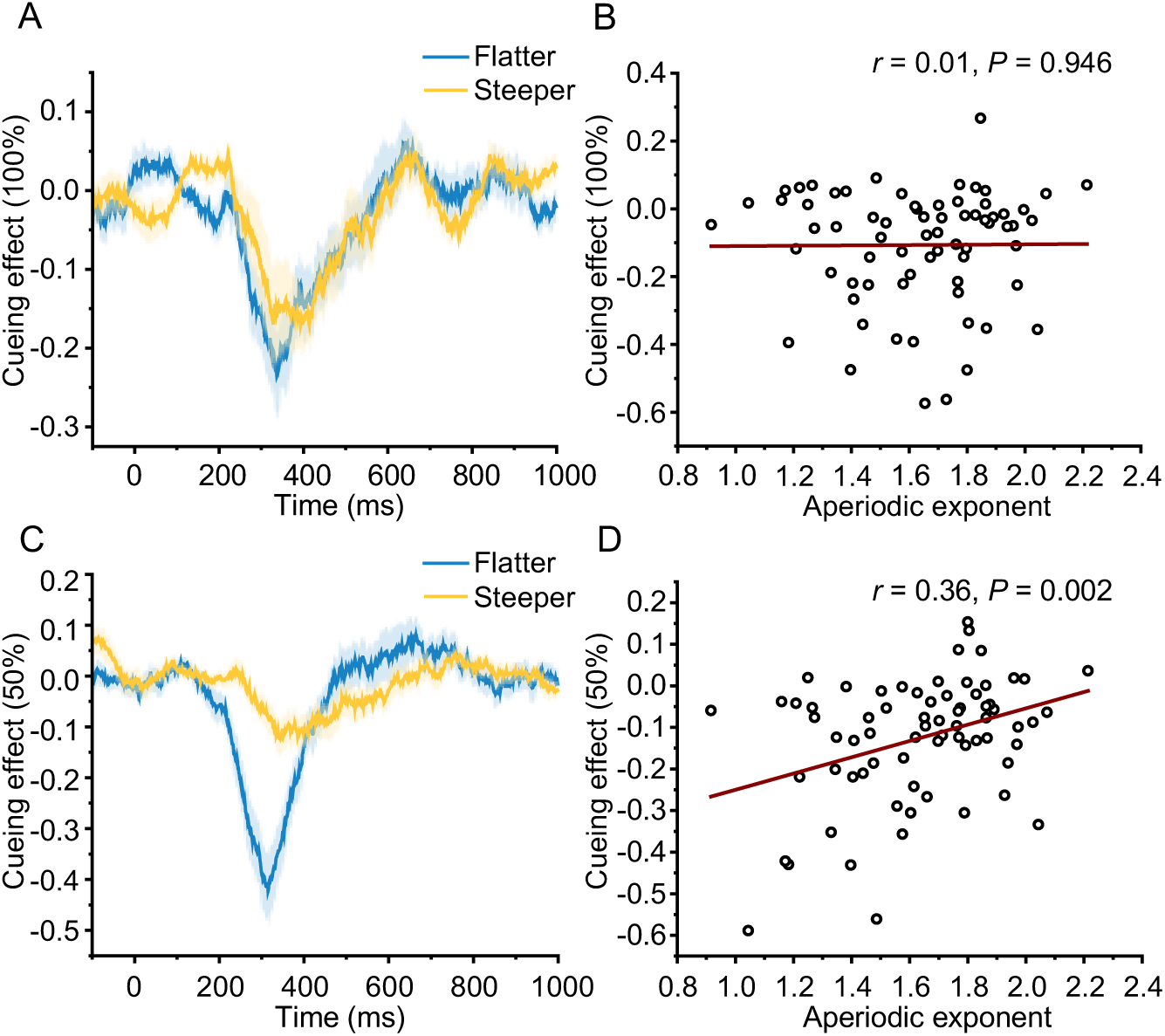
Relationship between the pupillary aperiodic exponent and the cueing effect in microsaccade rate. (A) Microsaccadic cueing effects for two groups of participants with flatter and steeper spectral slopes in the 100% cue-validity condition. (B) Correlation between the microsaccadic cueing effect and the aperiodic exponent in the 100% cue-validity condition. (C) Microsaccadic cueing effects for two groups of participants with flatter and steeper spectral slopes in the 50% cue-validity condition. (D) Correlation between the microsaccadic cueing effect and the aperiodic exponent in the 50% cue-validity condition. The shaded areas indicate the SEM.

For the 50% cue-validity condition, the cueing effect of microsaccade was also negative after cue presentation (Fig. 3C). Interestingly, the microsaccadic cueing effect exhibited an obvious difference between the two groups of participants, with the participants having lower aperiodic exponent (i.e., flatter spectrum) showing greater microsaccadic IOR. To validate the results of median splitting, we also extracted the averaged microsaccadic cueing effect in 200-500 ms for all participants and calculated its correlation with the aperiodic exponent of the pupil fluctuations (Fig. 3D). We found that the aperiodic exponent was significantly correlated with the microsaccadic cueing effect in the 50% cue-validity condition (*r* = 0.36, *P* = 0.002). The findings were consistent with the behavioral performance results, indicating that the aperiodic exponent was linked to both microsaccadic and behavioral responses only when the cue validity was 50%. These findings illustrate that individuals with a low aperiodic exponent showed increased sensitivity to nonpredictive cues and struggled to suppress automatic microsaccadic responses away from the cue. This suggests that the aperiodic exponent of resting-state pupil fluctuations may be indicative of the balance between excitatory and inhibitory processes in attentional functioning.

### Raw pupillary measurements did not predict microsaccadic IOR

To explore whether the prediction of the microsaccadic cueing effect in the 50% cue-validity condition was selective to the aperiodic exponent of pupil fluctuations, we further calculated the correlations between the microsaccadic cueing effect (mean value in 200-500 ms, as above) and the mean, variance, and hippus power of the raw pupil size. We found no significant correlation (all Ps > 0.482) between the microsaccadic cueing effect and any of these indices of raw pupil size (Fig. 4). This result suggests that the aperiodic exponent of pupil fluctuations predicts attentional processing better than the raw pupil size.

**Fig. 4.**
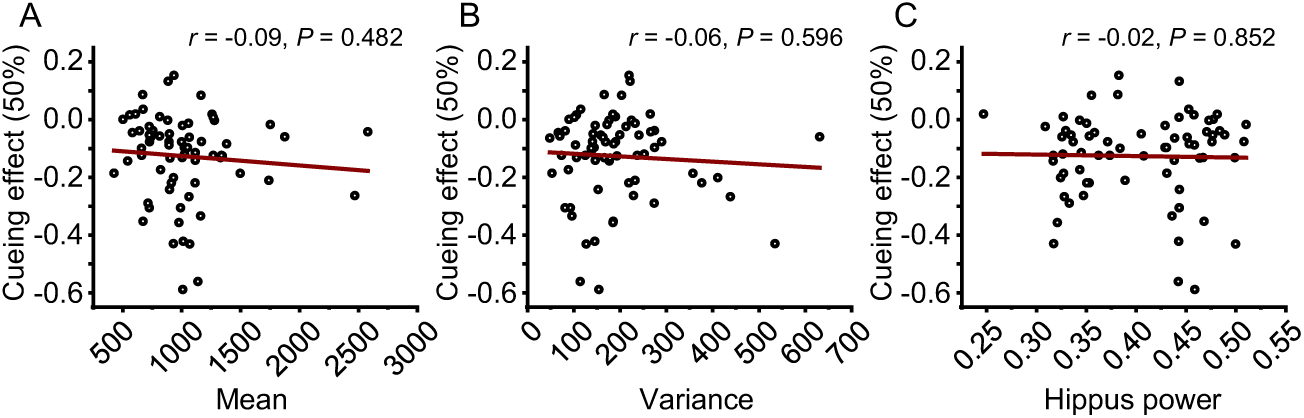
Correlations between the microsaccadic cueing effect in the 50% cue-validity condition and the mean (A), variance (B), and hippus power (C) of the pupil size data.

We found that the aperiodic exponent of pupil fluctuations was associated with cueing effects (both behavioral and microsaccadic) selectively in the 50% cue-validity condition, but not in the 100% cue-validity condition. This tentatively suggests that our analyses of the pupil data were valid. To further investigate whether the aperiodic exponent of the resting pupil fluctuations was affected by the interpolation of missing values (mainly caused by blinks) in the pupil data, we calculated the number of blinks during the resting state. We further examined the correlation between the aperiodic exponent and the number of blinks. The results showed no significant correlation (*r* = 0.02, *P* = 0.857). This further suggests that our analyses of the resting pupil data were valid and not affected by missing-value interpolation.

### Microsaccadic response mediated the effect of the aperiodic exponent of resting pupil fluctuations on the behavioral effect

Since the aperiodic exponent of pupil fluctuations was measured in the resting state and independent of the visual attention task, its correlation with microsaccadic IOR and behavioral effect in the 50% cue-validity condition indicates that the aperiodic exponent of resting pupil fluctuations may serve as an individualized trait of attentional processing. If so, then how would the pupil fluctuation influence the behavioral response to visual stimuli? We hypothesize that the pupil fluctuations may directly influence the microsaccadic IOR to the stimulus, which then influences the behavioral responses to the stimulus. Supporting this hypothesis, participant-wise mediation analyses revealed that the relationship between the aperiodic exponent of resting pupil fluctuations and the behavioral cueing effect was fully explained by the microsaccadic IOR (Fig. 5; indirect effect: *b* = 0.02, *P* = 0.043; direct effect: *b* = 0.03, *P* = 0.166; total effect: *b* = 0.05, *P* = 0.018). The total and indirect effects were significant in the mediation, but the direct effect was not significant. These outcomes indicate that the aperiodic exponent of resting pupil fluctuations influences individuals’ microsaccadic responses to the cue which then influence the behavioral performance in spatial orienting. These results support the hypothesis that the aperiodic exponent of resting pupil fluctuations serves as an individualized trait that affects behavioral outcomes by influencing microsaccadic responses.

**Fig. 5.**
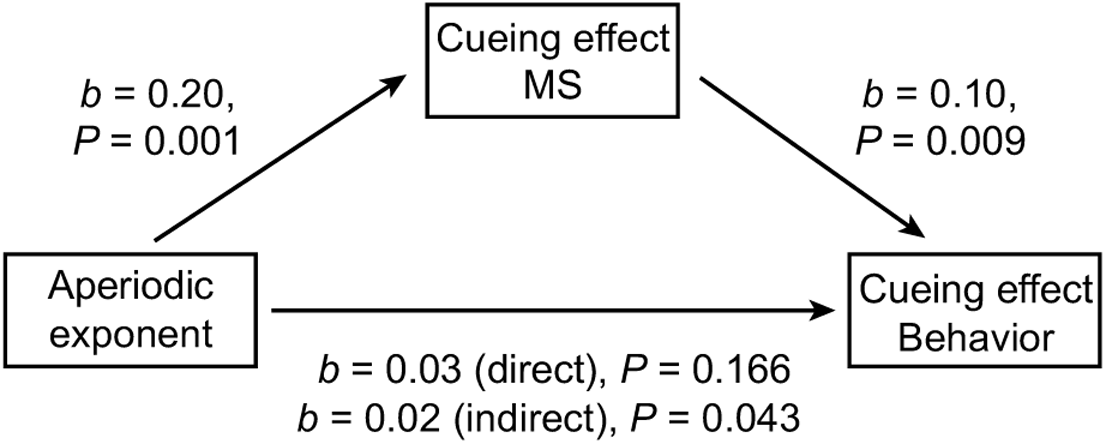
Participant-wise mediation model for pupillary and behavioral data in the 50% cue-validity condition. The cueing effect of microsaccade mediates the relationship between the aperiodic exponent and the cueing effect of behavior.

Given that the mediation analysis showed that the effect of the aperiodic exponent was explained through the microsaccadic cueing effect, there was a possibility that the individual difference in the aperiodic exponent indicated trivial differences in the quality of the eye-tracking recordings that also affected the analysis of the microsaccade. To investigate this possibility, we calculated the correlation between the aperiodic exponent of pupil fluctuation and the averaged microsaccade rates in the 1 s after cue onset in the 100% and 50% cue-validity conditions. The results showed no significant correlation in either the 100% (*r* = 0.14, *P* = 0.239) or the 50% (*r* = 0.17, *P* = 0.157) cue-validity condition. This suggests that the difference in the aperiodic exponent of pupil fluctuation did not affect the quality of the eye-tracking recordings, which would otherwise affect the overall detection rate of microsaccade.

### The aperiodic exponent of resting pupil fluctuations predicted self-rated ADHD-like symptoms

To further investigate the association between the aperiodic exponent of pupil fluctuations and the trait of attentional processing, we calculated correlations between the aperiodic exponent and the self-rated ADHD symptoms measured by ASRS (Kessler et al., 2005). We calculated the correlation between the aperiodic exponent and the total self-rated ADHD score as well as its inattention and hyperactivity/impulsivity subcomponents. We found a significant negative correlation between the aperiodic exponent and the total self-rated ADHD score (Fig. 6A; *r* = -0.27, *P* = 0.023), suggesting that a lower aperiodic exponent (flatter slope) was accompanied by more severe self-rated ADHD symptoms. Further analysis showed that the aperiodic exponent was significantly correlated with self-rated hyperactivity/impulsivity (Fig. 6B; *r* = -0.35, *P* = 0.003) but not with self-rated inattention (Fig. 6C; *r* = -0.10, *P* = 0.472). Therefore, a lower aperiodic exponent was accompanied by more severe self-rated hyperactivity/impulsivity. These results suggest that the aperiodic exponent of pupil fluctuations, like the aperiodic exponent of the EEG signals (Karalunas et al., 2022), could predict the level of self-rated ADHD symptoms.

**Fig. 6.**
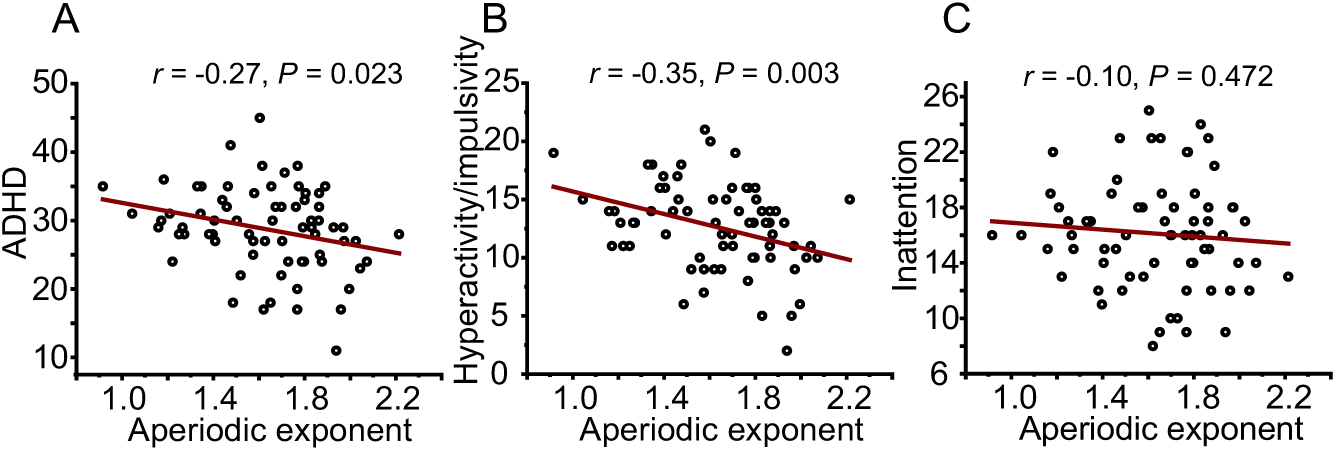
Correlations between aperiodic exponent and self-rated ADHD (A), self-rated hyperactivity/impulsivity (B), and self-rated inattention (C) symptoms.

As a control analysis, we also calculated the correlation between the total self-rated ADHD score and the mean, variance, and hippus power of the raw pupil size data. No significant correlations were observed in any of the analyses (Fig. 7). This result further confirmed that the aperiodic exponent was more predictive of the trait of attentional processing than other raw indices of pupil size.

**Fig. 7.**
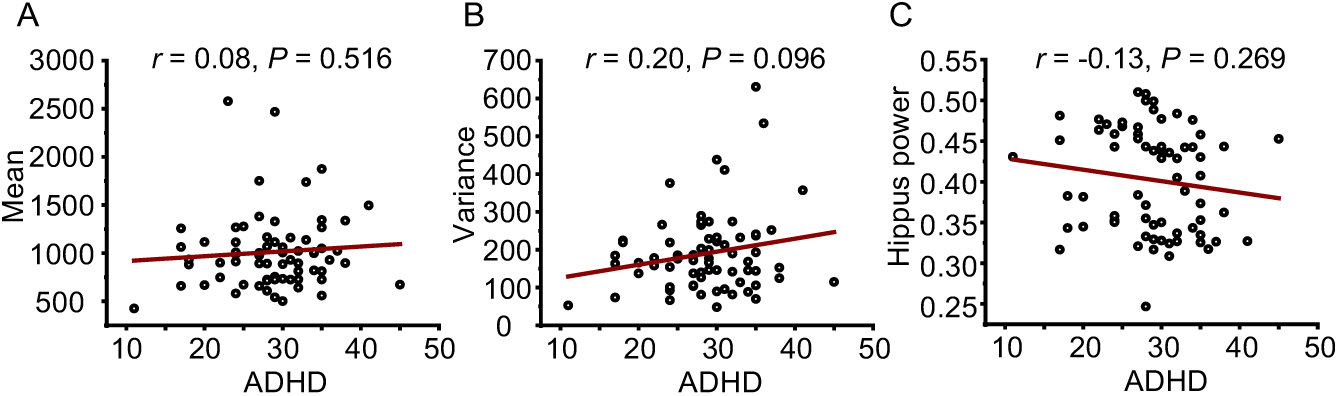
Correlations between the total self-rated ADHD score and mean (A), variance (B), and hippus power (C) of pupil size.

## Discussion

The present study investigated whether the aperiodic exponent of resting pupil fluctuations predicts neural E/I balance of attentional processing. We analyzed participants’ aperiodic exponent of pupil fluctuations in the resting state, examined their neural E/I balance with an attentional orienting task, and estimated their attentional traits with a self-rated ADHD scale. The aperiodic exponent of pupil size fluctuations correlated with the behavioral and microsaccadic cueing effects when the cue validity was 50%. Specifically, a greater aperiodic exponent was accompanied by a larger cueing effect of behavior and a lower microsaccadic IOR in the 50% cue-validity condition but not in the 100% cue-validity condition. A mediation model further identified that the microsaccadic IOR mediated the effect of the aperiodic exponent on the behavioral cueing effect. Finally, participants with a lower self-rated hyperactivity/impulsivity symptom exhibited a greater aperiodic exponent in their pupil fluctuations. Notably, because the pupil size information was recorded offline, these correlations were free from contaminations due to simultaneous eye movement or behavioral response recordings. Control analyses further showed that the prediction of the attentional effects and self-rated ADHD symptoms did not exist in other raw pupil size indices. Taken together, the aperiodic fluctuation of pupil size predicts neural E/I balance in attentional processing.

Instead of focusing on the raw pupil size data, the present study analyzed the aperiodic exponent of pupil fluctuations during the resting state and explored its potential functions in predicting neural E/I balance of attentional processing. This study is inspired by the widely demonstrated aperiodic exponents in neural signals and behavioral responses. For example, 1/f-like power spectrum has been revealed in EEG (Waschke et al., 2021), MEG (Dehghani et al., 2010), ECoG (Manning et al., 2009), and fMRI (He, 2011) data. Functionally significant aperiodic exponents have also been found in the analysis of human behavioral responses (Palva et al., 2013). The aperiodic exponent of neural signals has been shown to be related to the E/I balance in the neural system (Gao et al., 2017), with a greater aperiodic exponent, i.e., a steeper spectrum, indicating more neural inhibition while a lower aperiodic exponent, i.e., a flatter spectrum, indicating more neural excitation. The present study showed an association between the aperiodic exponent of resting pupil fluctuations and neural E/I balance in attentional processing. Specifically, the aperiodic exponent of pupil fluctuations was correlated with two different indices of spatial orienting, the microsaccadic responses and behavioral performance, in an attentional cueing task. Further, the aperiodic exponent of pupil size also predicted individuals’ self-rated ADHD symptoms of hyperactivity/impulsivity.

Excitation and inhibition are two fundamental processes that govern the flow of information in the brain. The balance between excitation and inhibition is critical for many brain functions, including attention (Deiber et al., 2020). Attention, defined as the ability to focus on a specific stimulus or task while ignoring others, is a selective process that allows us to filter out task-irrelevant information and concentrate on what is important. Thus, E/I balance plays a key role in attention by aiding in the selection of task-relevant stimuli for processing (Beesley et al., 2015). Additionally, E/I balance is also important for maintaining attention over time. Initially, when we attend to a stimulus, the E/I balance is tipped in favor of excitation, enabling quick and efficient processing of the stimulus. However, over time, the balance gradually shifts back towards inhibition. This prevents our attention from being depleted on one stimulus, providing the opportunity to shift focus to other potential targets in the environment (Jia et al., 2019; Lv et al., 2022). In the present attention tasks, participants must suppress the automatic microsaccadic responses toward exogenous cues to maintain fixation after cue onset (Rolfs et al., 2004). Consequently, the automatic microsaccades towards the cue are suppressed, while the microsaccades opposite to the cue are enhanced. The response properties of microsaccades reflect the balance of neural excitation and inhibition in participants during the attentional tasks. As shown in our results, participants with a lower aperiodic exponent exhibited over-excitation to the exogenous cue and over-suppression of the automatic microsaccadic response to the cue. We further demonstrated the mechanism underlying this association through a mediation model. We found that the association between the aperiodic exponent and the behavioral performance in the attention task was fully mediated by the microsaccadic responses. This may suggest that variations in neural E/I balance lead to different patterns of automatic microsaccadic responses to the cue, which in turn result in different attentional effects in behavior.

In the present results, the aperiodic exponent of resting pupil fluctuations was correlated with behavioral cueing effect and microsaccadic IOR in the 50% cue-validity condition but not in the 100% cue-validity condition. We speculate that this is because participants were certain of where the probe would appear in the 100% cue-validity condition, so they did not suppress the automatic microsaccadic response in the direction of the predictive cue. In contrast, in the 50% cue-validity condition, participants were completely uncertain about where the probe would appear. Thus, they tended to suppress the automatic microsaccadic response in the direction of a nonpredictive cue. Consequently, the neural E/I balance is critical for response control in the 50% cue-validity condition. These results replicate the previous findings that the E/I balance in attentional processing is sensitive to cue validity (Beesley et al., 2015) and stimulus uncertainty (Feldman & Friston, 2010; Yu & Dayan, 2005) and demonstrate that the aperiodic exponent of resting pupil fluctuations predicts the neural E/I balance in attentional processing.

The predictive effect of the aperiodic exponent of pupil fluctuations cannot be accounted by the different levels of fatigue/vigilance. This is because fatigue/vigilance is a general state that should affect behavioral and microsaccadic responses in all tasks. However, our results showed that, in behavior, the aperiodic exponent of pupil fluctuations was only associated with the cueing effect in the 50% cue-validity condition and not with response accuracies. Furthermore, the aperiodic exponent of pupil fluctuations was only associated with the microsaccadic cueing effect in the 50% cue-validity condition but not in the 100% cue-validity condition. This indicates that the aperiodic exponent of pupil fluctuation reflects a specific mechanism of attentional processing, namely attentional orienting, rather than general fatigue/vigilance levels.

Previous studies have identified a strong relationship between the aperiodic exponent of neural signals and ADHD, albeit showing opposite trends across different age groups (Karalunas et al., 2022; Ostlund et al., 2021; Robertson et al., 2019). Despite numerous investigations linking various pupil indices during resting states, such as size and hippus, with ADHD in adults (Nobukawa et al., 2021), none have explored the aperiodic exponent of pupil fluctuations or its relationship with ADHD. For example, a recent study has focused solely on the raw pupil size data (Bellato et al., 2023) and failed to predict ADHD symptoms. Here, by extracting the aperiodic exponent of the pupil fluctuations, we revealed a significant association between the aperiodic exponent and the self-rated hyperactivity/impulsivity symptom, underscoring the functional significance of aperiodic dynamics in pupil size compared to raw pupil size (Binda & Lunghi, 2017). Notably, the pupil size data in the present study were recorded offline during a 300-s resting state. It, thus, provides a possibility that offline eye recording could provide important information about cognitive functions and associated traits. Given that E/I balance, as indexed by the aperiodic exponent of neural signals, has been associated with a variety of neurological disorders, including ASD (Bast et al., 2023), ADHD (Karalunas et al., 2022), and Parkinson’s disease (Wiest et al., 2023), our study suggests that the aperiodic exponent of pupil fluctuations has the potential to serve as an indicator for assessing these neurological disorders.

In conclusion, the present study shows that the aperiodic exponent of resting pupil fluctuation predicts the neural E/I balance in attentional processing. Neural E/I balance, a trait of brain information processing, influences the microsaccadic and behavioral responses in attentional orienting tasks. The present study provides a valuable index for noninvasive measurement of neural E/I balance and emphasizes that pupil fluctuations could provide rich information about cognitive processing.

## Author contributions

**Rongwei Wang**: Investigation, Formal analysis, Visualization, Writing.

**Jianrong Jia**: Conceptualization, Formal analysis, Visualization, Writing, Supervision.

## Acknowledgments

This work was supported by the National Natural Science Foundation of China (32371086) and the Zhejiang Provincial Natural Science Foundation of China (LY23C090001).

